# MMINT: a Metabolic Model Interactive Network Tool for the exploration and comparative visualisation of metabolic networks

**DOI:** 10.1101/2024.08.06.606923

**Authors:** Juan P. Molina Ortiz, Matthew J. Morgan, Amy M. Paten, Andrew C. Warden, Philip Kilby

**Affiliations:** CSIRO Environment, Canberra, ACT 2601, Australia; CSIRO Microbiomes for One Systems Health – Future Science Platform, Australia; CSIRO Advanced Engineering Biology – Future Science Platform, Australia; CSIRO Data61, Canberra, ACT 2601, Australia

## Abstract

Genome-scale metabolic models (GEMs) are essential tools in systems and synthetic biology, enabling the mathematical simulation of metabolic pathways encoded in genomes to predict phenotypes. The complexity of GEMs, however, can often limit the interpretation and comparison of their outputs. Here, we present MMINT (Metabolic Modelling Interactive Network Tool), designed to facilitate the exploration and comparison of metabolic networks. MMINT employs GEM networks and flux solutions derived from Constraint Based Analysis (e.g. Flux Balance Analysis) to create interactive visualizations. This tool allows for seamless toggling of source and target metabolites, network decluttering, enabling exploration and comparison of flux solutions by highlighting similarities and differences between metabolic states, which enhances the identification of mechanistic drivers of phenotypes. We demonstrate MMINT’s capabilities using the *Pyrococcus furiosus* GEM, showcasing its application in distinguishing the metabolic drivers of acetate- and ethanol-producing phenotypes. By providing an intuitive and responsive model-exploration experience, MMINT addresses the need for a tool that simplifies the interpretation of GEM outputs and supports the discovery of novel metabolic engineering strategies. MMINT is available at https://doi.org/10.6084/m9.figshare.26409328

**Graphical abstract:** *MMINT functionalities provide an intuitive and responsive model-exploration experience, enabling flux solution comparison and the identification of metabolic drivers of phenotypes*

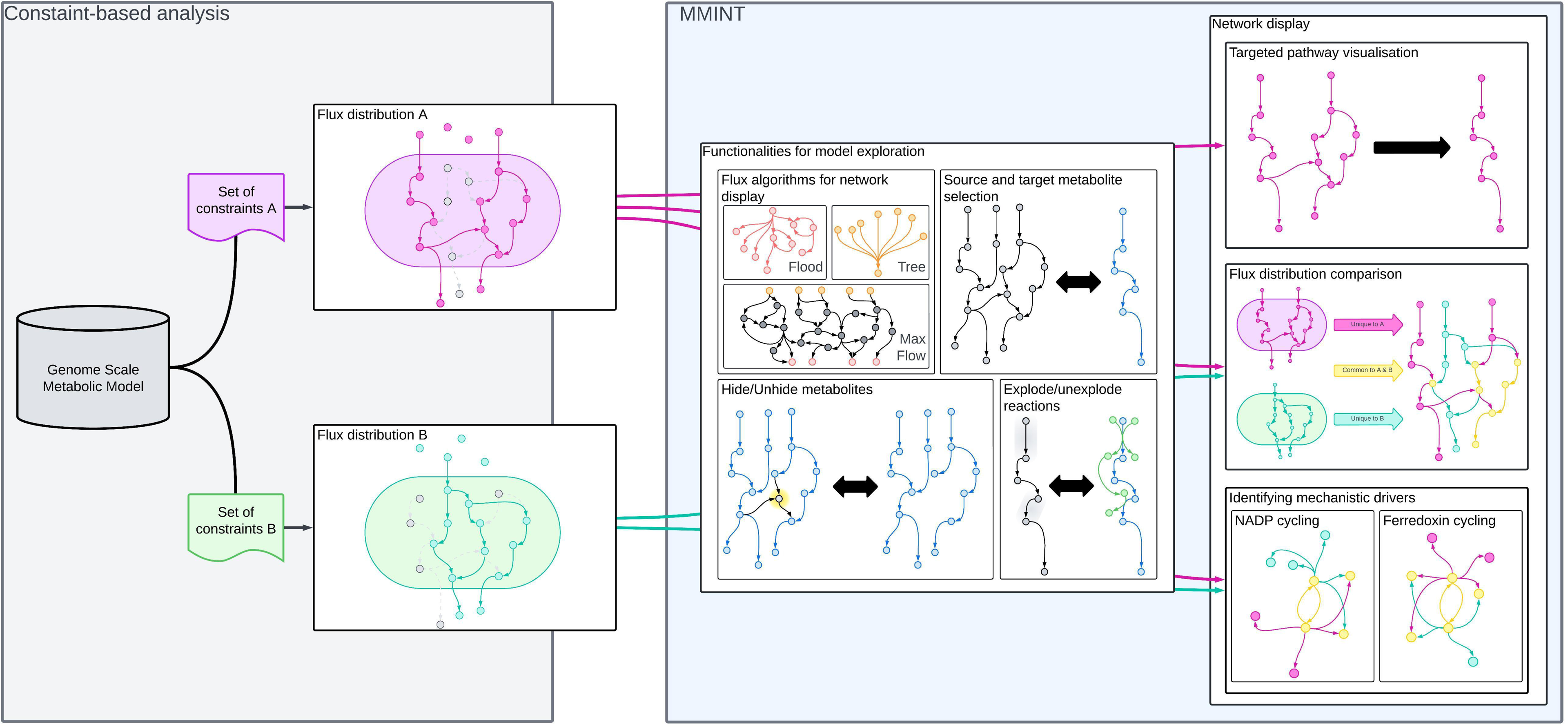

## Introduction

In the fields of systems biology and synthetic biology, genome-scale metabolic models (GEMs) have become instrumental tools for exploring and manipulating cellular metabolism. GEMs enable mathematical simulation of metabolic pathways, linking genes, proteins, and reactions through networks to predict metabolic phenotypes (1–3). Through the application of Constraint Based Analysis (CBA), such as Flux Balance Analysis (FBA) (4), GEMs can elucidate metabolic network adaptations in different cell types, organisms, and microbial communities under different environmental or genetic variations (5–7). By tracing flux distributions through a given metabolic network, GEMs-based analyses can identify metabolic mechanisms underlying emergent phenotypes (5,8). The complexity and mathematical format of GEMs, however, can make interpretation and comparison of outputs challenging and often limits the mechanistic understanding GEMs strive to provide.

The rapid advancement in GEM-building pipelines (9–14) and the availability of online databases with reliable reconstructions (15–19) have enabled the application of these models across numerous research areas, including enzyme engineering, human health, and microbial ecology (1,6,7,20–23). Underpinning their utility is the ability to explore and compare flux distributions. In synthetic biology, algorithms such as Optknock (24) and FastKnock (25) use GEMs to guide gene deletion or regulation strategies for optimizing bioproduction. Similarly, in microbial ecology, flux exploration aids in identifying cross-species metabolic interactions in microbial communities and host-microbiome systems. Tools such as MICOM (26) leverage GEMs for community metabolic flux analysis. For instance, Marcelino et al. (27) used MICOM and GEMs automatically reconstructed from prokaryotic metagenome-assembled genomes to investigate microbial community interactions in Crohn’s disease patients. This study uncovered metabolic mechanisms underlying the disease state, including a loss of cross-feeding. In another study, Schafer et al. (19) compared metabolic flux distributions to explore pairwise interactions in plant leaf microbiota as a framework for informing the construction of synthetic microbial communities according to substrate. Therefore, there is a need for tools that can rapidly assess and compare outputs from automatically inferred GEMs or models built by third parties, enabling researchers to determine their suitability for downstream applications. This underscores the need for a tool specifically designed for model and flux exploration and metabolic phenotype comparison.

Visualisation tools can be used to support the flux tracing process, addressing the complexity of interpreting GEMs solutions, and enabling comparison and examination of flux distributions. Yet, platforms such as Escher (28), KBase (29–31) and CAVE (Cloud-based platform for the Analysis and Visualisation of mEtabolic pathways) (32) require a comprehensive understanding of the network encoded in a GEM. For instance, Escher relies on a preexisting metabolic map built around the GEM in question to display fluxes from a given CBA solution. Meanwhile, CAVE relies on the manual selection of individual reactions to visualise them as a network. Further, these tools focus on individual model visualisation without offering an intuitive and direct comparison of alternative flux distributions, which can guide the identification of the mechanisms that drive the emergence of divergent phenotypes. Cytoscape (33), which has been adapted (e.g., as a plugin) to visualise metabolic networks, allows for the display of one or more CBA solutions. However, this process involves manually converting metabolites and reactions into network nodes and edges which can be time-consuming.

To address these limitations, we introduce MMINT (Metabolic Modelling Interactive Network Tool), a tool specifically designed for the exploration and comparison of metabolic networks. MMINT leverages GEM networks, which can be combined with up to two CBA solutions, to create an interactive representation of flux distributions through such network. When two solutions are provided, MMINT highlights their differences and similarities. MMINT provides a responsive model-exploration experience through network ‘decluttering’ functions, including seamless toggling of source and target metabolites, delivering immediate visual feedback, and enabling users to focus on specific pathways of interest. In doing so, these features further facilitate the identification of mechanistic drivers of phenotypes.

In this paper, we detail MMINT’s functionalities and applications using a pre-existing GEM of *Pyrococcus furiosus* (iPfu) (5), a thermophile archaeon and metabolic engineering platform. By comparing two engineering strategies derived from iPfu, we show how MMINT can contribute to model exploration and to the identification of metabolic phenotypic drivers. The potential impact of MMINT extends beyond this example. By enabling the interactive visualization of metabolic flux and interrogation of the quality of newly built or third-party GEMs, MMINT can become an indispensable tool for researchers aiming to gain deeper insights into metabolic networks and their applications in synthetic biology and microbial ecology.

## Implementation

### Software/Computational resources

The MMINT system is written in Java, using OpenJDK 22. MMINT requires the Java Run Time Environment (JavaRTE), which is available for all major operating systems. MMINT also relies on the “dot**”** executable from the *GraphViz* package (34). GraphViz is an open-source graph visualization package, available for all major operating systems. The dot program is used for graph layout by MMINT and can take a long time to process graphs with many edges and nodes. We limit the calculation time to 10 seconds. MMINT runs in a single thread on the local computer. All data is kept locally. Data privacy is ensured, as no data is sent to, or read from, remote servers. Once installed, MMINT can be used without access to the internet.

#### Input data and storage

Several types of input can be used for MMINT

- A GEM, describing reactions stoichiometries and gene-protein-reaction associations, in an XML/SBML format.
- Up to two CBA “solution” files. Solution files record the flux distribution through the reactions encompassed in a GEM during constraint-based analysis. Solution files should include reaction identifiers in the first field/column and the corresponding flux in the second field. Solution files can be provided in several formats.

`o` CSV (comma-separated values).
`o` TSV (tab-separated values)
`o` Text: one or more spaces separate the two fields.
`o` XLSX (Microsoft Excel file).

A ℌdrag-and-drop” interface is available to use files from the local system. All data is kept locally, in text-based (rather than binary) files.

#### Design and network visualisation

The MMINT software is composed of two interactive frames or “windows”. A control frame and a graph frame. The control frame handles file loading and lists the names of metabolites and reactions in a GEM and CBA solution. Meanwhile, the graph frame displays a graphic representation of the fluxes (edges) in a CBA solution using nodes that represent metabolites and reactions in a GEM. Reactions are displayed in rectangles and metabolites in ellipses. Different colours and line style signify characteristics of the node - for example gene-indicated reactions are displayed in rectangles with vertical double lines (see Figure S2 in Supplementary file 5 for legend of all variations of node characteristics). Importantly, MMINT does not display zero-flux reactions (non-active). Edges into a reaction indicate reactants and edges out represent products of the reaction. Hovering the mouse over reactions, metabolites, and edges displays stoichiometric information, formulas, and flux quantities, respectively.

MMINT uses an automatic graph layout to organise network elements displayed in the graph frame. This means that modifications to the displayed network can be quickly incorporated, with no setup required – as soon as a GEM file and a CBA solution file are supplied, a graph is displayed.

### Main Functionalities

Full flux networks in CBA solutions can be highly complex, encompassing potentially hundreds or thousands of reactions and metabolites. Therefore, only a portion of the non-zero flux reactions (active), and the corresponding metabolites, are shown by default in MMINT. The tool relies on different functions that allow refinement and further exploration of networks shown by default.

### Media and product

The concepts of “media” and “product” define which elements of the network are displayed in the graph frame by default. Media are metabolites that are represented in the initial growth media registered in a CBA solution (products of active exchange reactions) and constitute the starting point of the metabolic process. Products are a) the reactants of the biomass growth reaction, and b) metabolic end products (reactants of active exchange reactions). Biomass reactant metabolites are highlighted because the production of these metabolites constrains the growth of the organism. General “intermediate” metabolites linking those from media and product are also displayed in the graph frame by default. Lists of media and product elements are shown in the MMINT control frame, which can be selected or unselected to modify the displayed network in the graph frame. Similarly, metabolites in general can be added to the media and/or product list in the control frame by right-clicking on the corresponding nodes on the graph frame and selecting “Add to –”.

### Algorithms

The default metabolic network shown in MMINT is also influenced by display algorithms. When exploring a flux distribution solution, different representations of the data can help to clarify the flow of elements through the system. For that purpose, MMINT provides several different algorithms to display the data; these can be selected from the *Algorithms* menu in the control frame.

We use a graph *G(N,E)* with nodes *N = M U R,* and edges *E*. *M* is the set of nodes representing metabolites (this includes media and product metabolites), and *R* is the set of nodes representing reactions. The set of edges in *E* is made of two types: edges in *M x R* from nodes representing the reactants to the reaction node itself; and edges in *R x M* from the reaction node to each of the products of the reaction.

Nodes with in-degree zero in *G* are called *source* or *media* nodes i.e., metabolites available in the media. Nodes with out-degree zero are included in the *product* nodes, and represent metabolites produced by the organism during growth (for example CO2). Product nodes also include reactants of the biomass reaction.

#### Max Flow algorithms

The **Max Flow** algorithm solves the classical Max Flow problem from graph theory (35), using an implementation of the Ford-Fulkerson algorithm (35). All edges in the graph have capacity equal to the flux through the associated reaction. As usual, with multi-media and multi-product instances, two dummy nodes *S* and *T* are added to *G*. Edges are added from *S* to all media nodes, and from all destination nodes to *T*. The maximum flow is then found between *S* and *T*. In terms of network visualisation, the Max Flow algorithm tries to find as many paths as possible between media metabolites and product metabolites. The paths are limited by the same flux as in the solution. Essentially, it will display all non-looped paths between selected media and products.

The **Simple Max Flow** algorithm solves the same problem on the same graph as Max Flow but uses capacity one on all edges. This translates into a visually simpler solution that shows only one pathway linking media and product metabolites. This is the default algorithm in MMINT.

#### Shortest Path algorithms

The **Shortest Path** and **Tree** algorithms use the Dijkstra algorithm (36) to find the shortest path from every media node to every destination node. Tree algorithms grow a tree of pathways from the selected media or product metabolite(s). Source trees use the shortest path from each selected media node to all selected destination nodes. Destination trees use the shortest path from selected media nodes to each selected destination. A media tree for instance, grows a tree from selected media metabolite(s), and shows all the reactions and metabolites that this media has a part in producing. The product tree is the opposite: it grows the tree *towards* the selected metabolite(s), showing all the reactions and metabolites that contribute to producing the selected product.

#### Flood algorithms

**Flood** algorithms also use the Dijkstra algorithm with a limited depth *D*. The length of every edge is set to one. Every edge from *G* expanded by the Dijkstra algorithm is added to the display graph. If an edge returns to a previously seen node, the edge is marked as a cycle edge, and the destination node is not considered again. Similarly, a node is not considered if the distance to the node exceeds *D*. In MMINT, flood algorithms find all possible paths from or to a metabolite. Since there are many such paths, the number of edges in the path is limited using the “Level” counter on the graph frame. Click “+” to increase the number of edges displayed, or “-” to decrease it.

#### Algorithm menu in MMINT

- Highlight flood: flood to and from the highlighted reaction or metabolite. Use “Level” buttons to adjust depth.
- Highlight tree: grow a media tree from, and also a product tree towards, the highlighted metabolite or reaction.
- Media or Product Flood: flood *from* all selected media metabolites, and *to* all selected product metabolites simultaneously. This option works best with a single media and single product and shows all possible ways to get from media to product metabolites. Use “Level” buttons to adjust depth.
- Media Tree: grow a tree from all selected media metabolites. Shows which metabolites or reactions the selected media metabolites produce, directly or indirectly.
- Media Flood: flood from all selected media metabolites. Shows every possible connection from the media metabolites. Use “Level” buttons to adjust depth.
- Product Tree: grow a tree towards all selected products. Shows which metabolites or reactions are directly or indirectly involved in the production of the selected product metabolites.
- Product Flood: flood towards all selected media metabolites. Shows every possible connection to the products. Use “Level” buttons to adjust depth.
- Max Flow: the max flow algorithm (described above)
- Simple Max Flow: the simple path max flow algorithm (described above)
- Shortest Path: find the shortest path between every pair of selected media and product metabolites. Only one path is shown, so some metabolites might not be shown if there is another path of the same or longer length between media and product.
- Longest Path: opposite to shortest path - finds the longest (non-looped) path between selected media and product metabolites.

### Additional functions

*Media*, *Product* and *Algorithms* have a broad effect over the network displayed in the graph frame. However, targeted modifications are often necessary when exploring metabolic networks and CBA solutions. The following MMINT functions enable control over elements displayed at an elemental level.

- Find element: a search function to identify a specific metabolite or reaction among displayed elements (reactions and/or metabolites).
- Hide metabolite: removes the selected metabolite from the displayed network.
- Explode or Unexplode element: reveals or hides, respectively, all the edges linked to the selected element to show additional pathways.

Together, these functions allow granular GEM and flux distribution exploration. The following section shows how they work in synergy to achieve specific tasks.

### Case Studies

To showcase MMINT’s capabilities, we introduce three distinct case studies centred around two *Pyrococcous furiosus* GEM (iPfu) phenotypes; one that overproduces acetate (acetate phenotype) and one that produces ethanol instead (ethanol phenotype) (5).

- Case one shows how the default network generated by MMINT can be refined to allow tracing of carbon fluxes to the Biomass reaction enhancing our understanding of pathway/reaction utilisation while hiding less relevant fluxes.
- Case two focuses on highlighting the differences in the metabolic networks in the acetate and ethanol iPfu phenotypes shedding light on the distinct metabolic adjustments that underpin each phenotype.
- Case three examines the underlying metabolic drivers responsible for the emergence of these phenotypes.

Together, these case studies exemplify how MMINT can be effectively employed to facilitate GEM exploration and understanding of the mechanisms from which different phenotypes emerge.

#### The *Pyrococcus furiosus* GEM (iPfu)

Vailionis *et al.* (5) built iPfu from GenBank assembly ASM730v1, which corresponds to *P. furiosus* DSM 3638. The model was manually curated and calibrated based on experimental growth data with three carbon sources (37), achieving a coefficient of determination (R²) of 0.95. The base iPfu model was later modified to resemble COM1, a *P. furiosus* strain amenable to genetic engineering. This process required removal of 15 metabolic reactions, with no reported impact to iPfu-derived growth prediction accuracy. The final characteristics of iPfu are shown in Table 1.

**Table 1.**
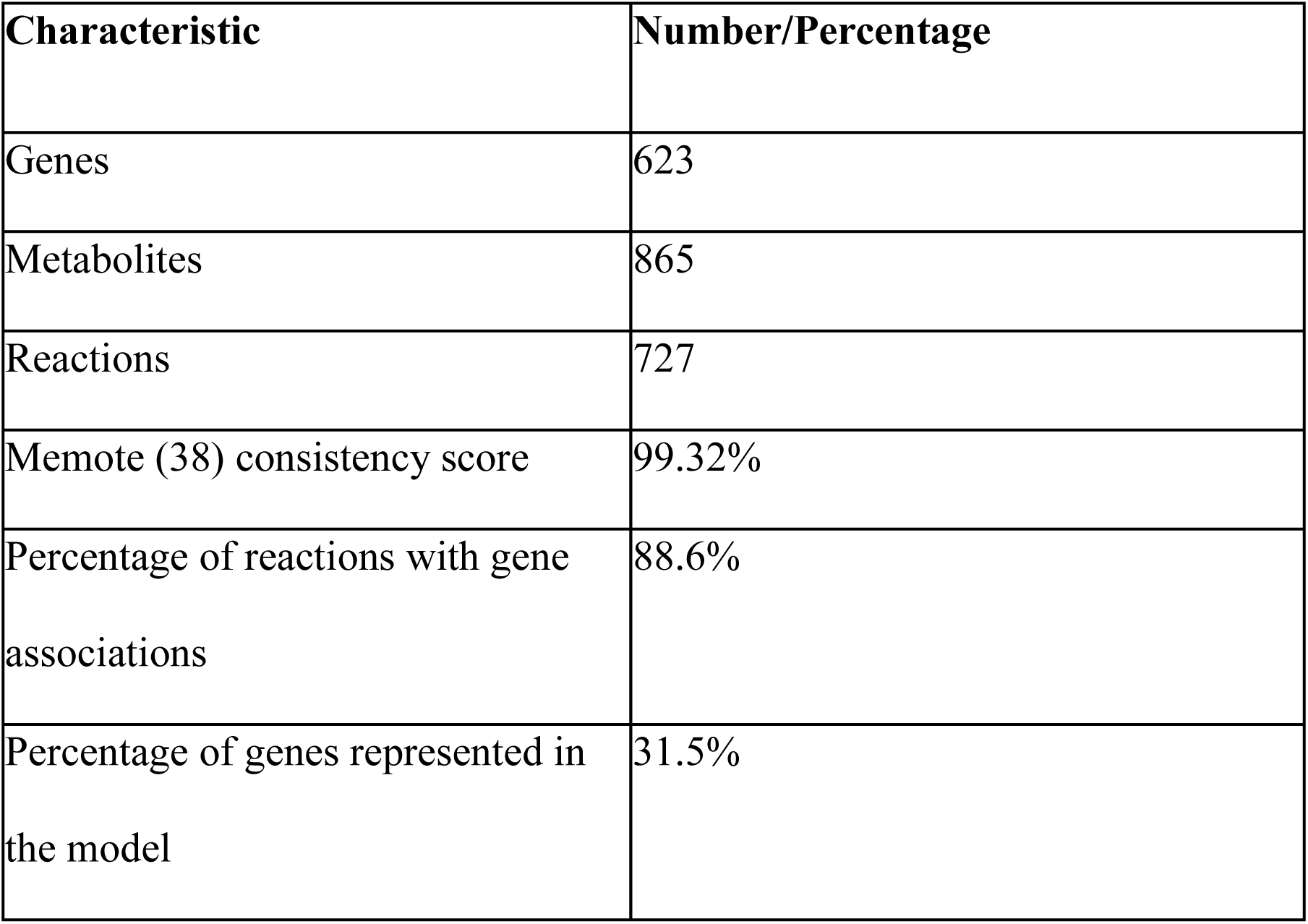
Characteristics and metrics of iPfu.

### iPfu variants

#### *OE-AdhF* (acetate phenotype)

Following extensive testing, an iPfu variant, OE-AdhF, was selected as a base model to explore potential engineering strategies aimed at optimising ethanol production in 14.6 mM maltose and 0.5 g/L yeast extract enriched media. OE-AdhF represents a variant where gene *adhF*, encoding an aldehyde dehydrogenase is over expressed, resulting in an increase in ethanol production. Gene overexpression is not explicitly captured in conventional GEMs such as iPfu. Therefore, the base model was constrained to produce a predetermined quantity of ethanol. Specifically, the lower bound of the ethanol transport reaction (TP_ETOH) was set to 3.29 mmol/gDW/h (millimoles per gram of dry weight per hour), which was determined from empirical data and random flux sampling analysis (5).

Optimising model OE-AdhF for biomass production in the specified media results in the synthesis of 3.29 mmol/gDW/h of ethanol, 46.09 mmol/gDW/h of acetate and 0.71 gDW/h of biomass (Figure 1A).

**Figure 1.**
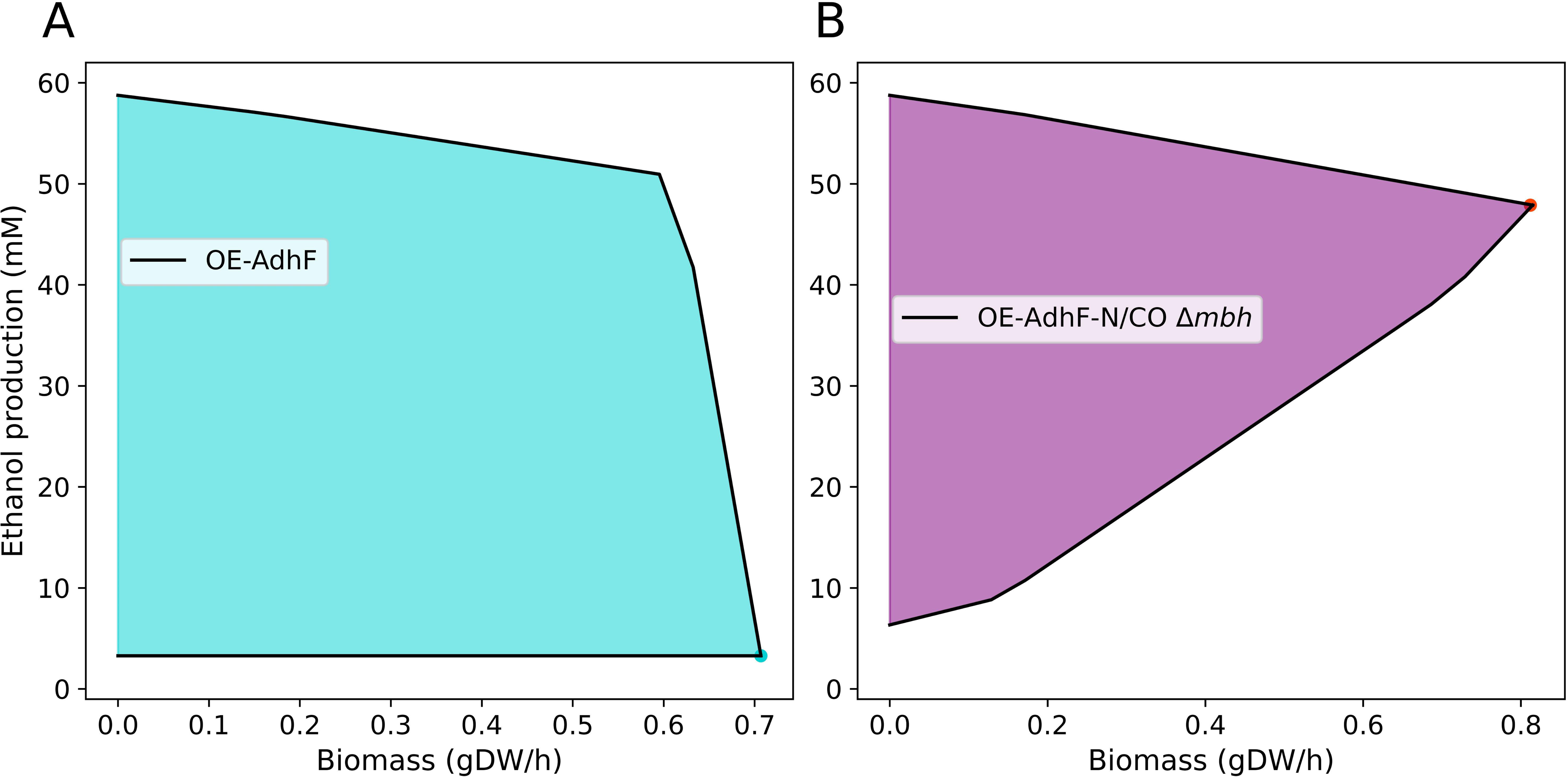
Ethanol-biomass production envelopes in OE-AdhF (acetate phenotype, light blue) and OE-AdhF-N/CO Δmbh (ethanol phenotype, purple). A) Ethanol production remains at an unchanged minimum (lower constraint of 3.29 mmol/gDW/h) regardless of changes in biomass. B) Minimum predicted ethanol production is directly proportional to biomass production. For our case-examples we rely on the flux distributions each variant achieves while maximising biomass production, highlighted with light blue and purple dots in plots A and B for the acetate and ethanol phenotypes, respectively.

Henceforth, we refer to this variant as the acetate phenotype.

#### *EO-AdhF-N/CO Δmbh* (ethanol phenotype)

Aiming to identify engineering strategies that favour ethanol production over acetate, Vailionis *et al.* sampled a considerable combination of gene insertions and deletions for OE-AdhF. The variant that required the least modifications and produces the most promising predictions is OE-AdhF-N/CO Δ*mbh*, which required two gene insertions and one deletion (Table 2). This variant, henceforth referred as the ethanol phenotype, is predicted to produce ethanol at a rate of 47.9 mmol/gDW/h, no acetate, and 0.8 gDW/h biomass (Figure 1B).

**Table 2.** Sequential modifications in OE-AdhF that lead to OE-AdhF-N/CO Δmbh. . The variant corresponding to the acetate phenotype is shown in light blue and the variant corresponding to the ethanol phenotype is highlighted in purple. *Variant name equivalent to model b971180d_YM_GAPN_CODH_closed_OEadhf.

| Variant name | Parent model/strain | Modification |
| --- | --- | --- |
| OE-AdhF | COM-1 | Strain: overexpression of adhF, which encodes an aldehyde dehydrogenase.<br>Model: lower bound of TP_ETOH (ethanol transport) constrained to 3.29 mmols/gDW/h. |
| OE-AdhF-N | OE-AdhF | Insertion of reaction GAPN, which converts glyceraldehyde-3-phosphate (GAP) into glycerol-3-phosphate (G3P) with NADP as intermediary. |
| OE-AdhF-N/CO* | OE-AdhF-N | Insertion of reaction CODH, which converts CO into CO<sub>2</sub>, reduces hydrogen into H<sub>2</sub> and exports sodium to the extracellular space. |
| OE-AdhF-N/CO Δmbh | OE-AdhF-N/CO | Deletion of mbh, which oxidises ferredoxin and reduces hydrogen into H<sub>2</sub>, while exporting sodium to the extracellular space. |

### Flux balance analysis (FBA) in iPfu variants

To replicate the metabolic states shown in Figure 1, we utilised the models b971180d_YM_closed_OEadhf (acetate phenotype) and b971180d_YM_GAPN_CODH_closed_OEadhf in their SBML format, which are part of (5) data repository (Supplementary file 1). Model b971180d_YM_GAPN_CODH_closed_OEadhf was further modified to represent the required ethanol phenotype (Table 2). To aid interpretability of the figures shown in the case studies below, relevant metabolite identifiers in model b971180d_YM_GAPN_CODH_closed_OEadhf were replaced with their corresponding metabolite names (Supplementary file 1 and 2). Compartment suffixes were also modified to represent metabolites in the extracellular space (_e) or intracellular space (_c). FBA (4) for each model was performed on the COBRAPy platform and the Gurobi solver (39) with biomass set as optimisation objective (40). FBA solutions were converted to Pandas DataFrame objects and exported as Comma Separated Value (.csv) files (Supplementary files 3 and 4). A detailed guide to replicate the networks shown in Case Studies (Figures 2, 3 and 4, respectively) can be found in Supplementary file 5. Supplementary file 6 contains the files required to replicate the flux distributions for the acetate and ethanol phenotypes.

**Figure 2.**
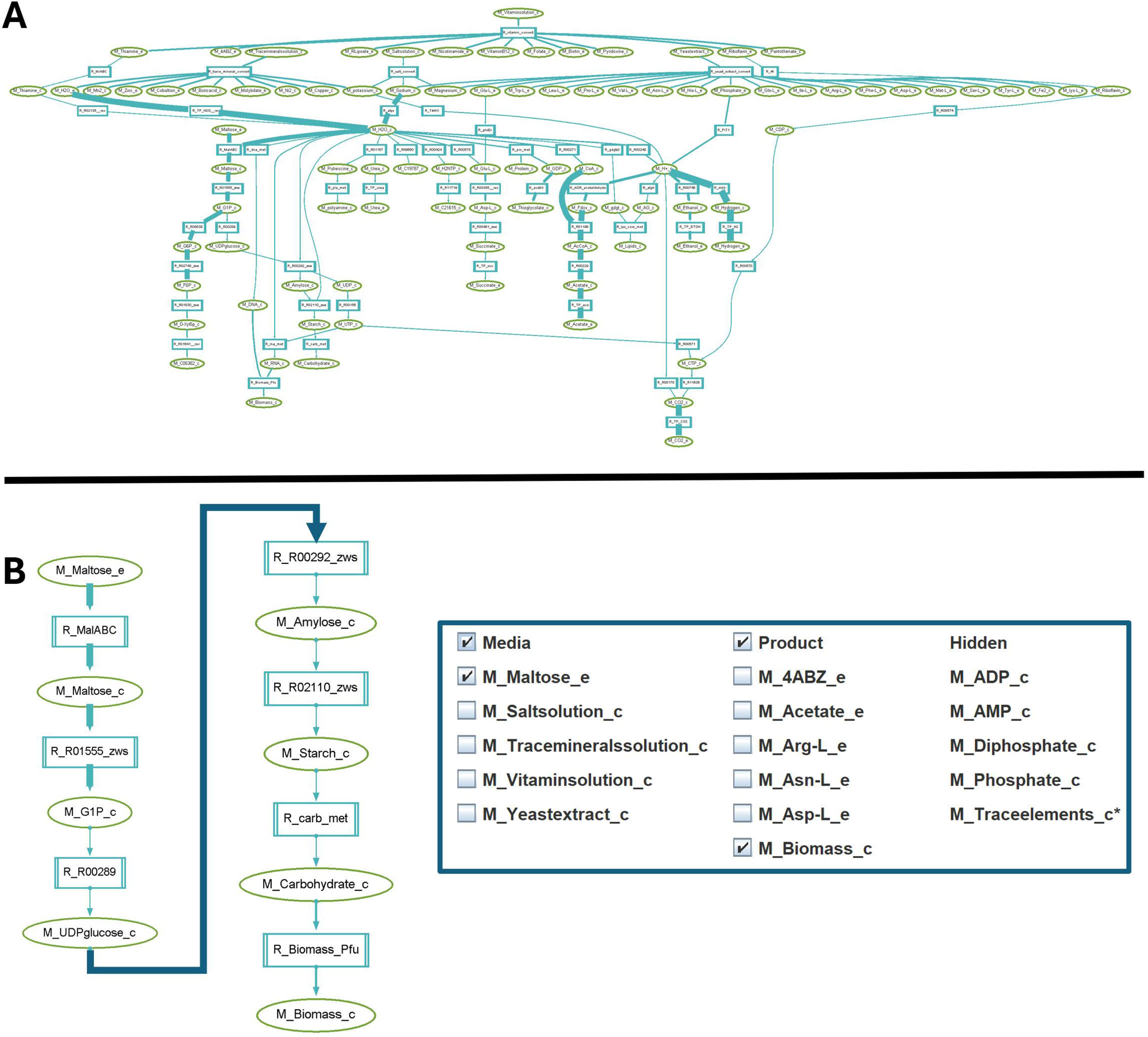
Targeted pathway visualisation. MMINT allows users to identify and display metabolites, reactions and pathways of interest while hiding less relevant information. A) Default network generated by MMINT, based on the fluxes derived from FBA in the acetate phenotype, which can be modified to show network elements of interest. B) Refined network generated by MMINT, from the original default network view, showing fluxes that connect maltose metabolism (M_Maltose_e) to biomass generation (M_Biomass_c). A capture of MMINT’s interactive panel shows metabolites chosen as media and product, and hidden metabolites, that lead to the network configuration shown in panel B. Metabolites are shown in light green and reactions are shown in light blue. Pseudo-metabolites shown under Media (Saltsolution, Tracemineralssolution, Vitaminsolution, Yeastextract) functionally work as extracellular metabolites.

**Figure 3.**
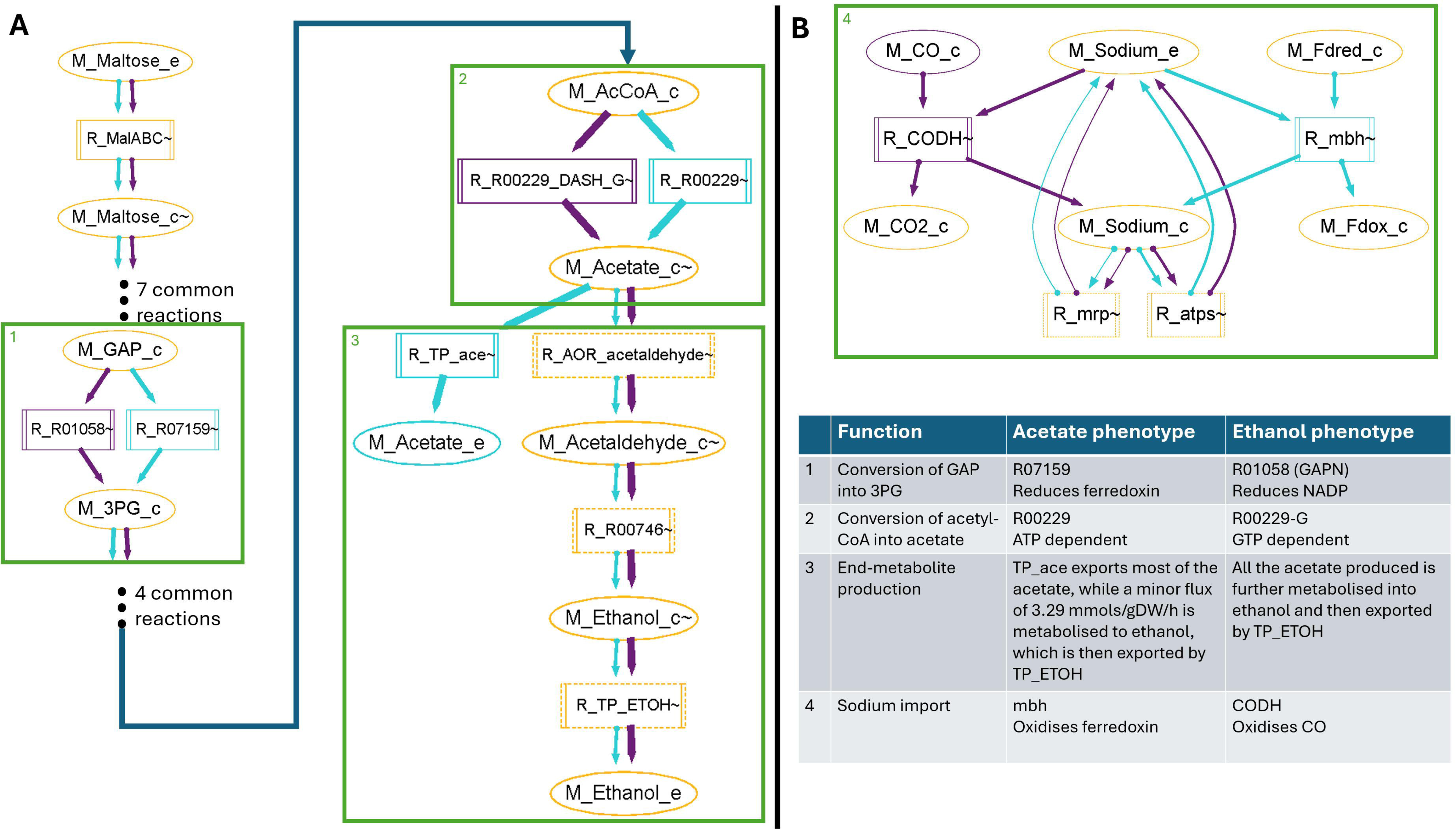
Flux distribution comparison. A) Differences in maltose degradation fluxes leading to the production of main end- metabolites between the acetate phenotype (light blue network elements) and the ethanol phenotype (purple network elements). Reactions and metabolites common to both phenotypes are shown in gold. A full display of the carbon fluxes in the Acetate and Ethanol phenotypes, including sections in common, is shown in Figure S 1 (Supplementary file 5). B) Differences in sodium import fluxes (Sodium_e to Sodium_c) resulting from knocked and inserted reactions in the ethanol phenotype in relation to the acetate phenotype. While most of the reactions that drive this process are common to both phenotypes, metabolic differences are displayed (green boxes and numbers). Table summarising main metabolic differences between the acetate and ethanol phenotype and the functions and reactions involved is included in the figure.

**Figure 4.**
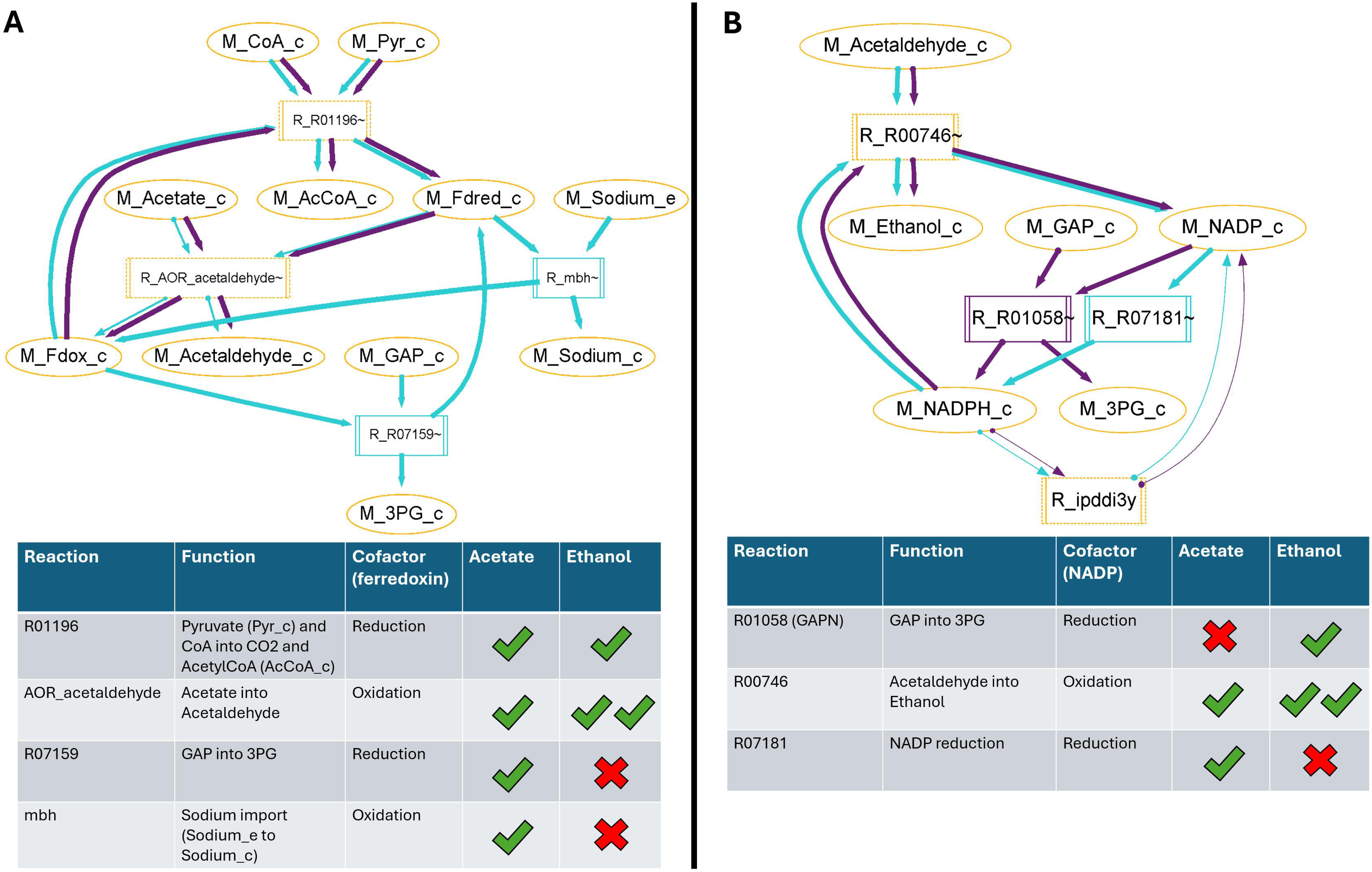
Cofactor cycling in the acetate and ethanol phenotypes. A) Main reactions and fluxes involved in cycling ferredoxin in the acetate and ethanol phenotypes. B) Main reactions and fluxes involved in cycling NADP in the acetate (light blue network elements) and ethanol phenotypes (purple network elements). Reactions and metabolites common to both phenotypes are shown in gold. Both networks shown in A and B were generated using functionalities in MMINT and are accompanied by a table with information about reactions shown in each representation. Zero-flux reactions in a given phenotype are indicated with a red cross, while reactions with non-zero fluxes are shown with a green check mark. Two check marks show reactions in a phenotype with increased fluxes with respect to the alternative phenotype.

### Case 1: Targeted pathway visualisation

After loading a model and a solution, MMINT will automatically identify media (inferred from extracellular metabolites in media) and product metabolites (inferred from output metabolites) and will build a default network that incorporates non-zero flux reactions. While such a network may already provide meaningful insights in terms of GEM structure and flux distributions, further refinement might be needed to identify metabolites, reactions, and pathways of interest. Figure 2 shows how combining various functionalities in MMINT allows users to rapidly identify such relevant elements and hide others. The main carbon source in iPfu default media is maltose, and in this example, we identify how maltose (Maltose_e) import and metabolism leads to biomass production (Biomass_c) in the acetate phenotype solution. This is achieved by selecting the specific metabolites as Media and Product, correspondingly, and hiding common cofactors and/or metabolites. Hiding metabolites that are common to many reactions is recommended as this will facilitate identification of more specific products and reactants.

### Case 2: Solution comparison

MMINT has been specifically designed to visualise and compare fluxes from two flux distribution solutions. Here, MMINT is used to focus on the similarities and differences in the fluxes that derive from maltose metabolism, and lead to the production of acetate and ethanol, in each of the corresponding phenotypes. To generate the network visualisations shown in this section, we rely on several MMINT functionalities, including Media and Product selection, hide/unhide and the explode and unexplode functions. Figure 3A shows that both phenotypes rely on a remarkably similar pathways but with notable differences, which are partially explained by reaction insertions and deletions.

First, the acetate phenotype utilises R07159 to metabolise GAP into 3PG, while the ethanol phenotype relies on R01058 (GAPN) for the same metabolic conversion (Figure 3A, box 1). A crucial difference between these two reactions is that GAPN reduces NADP as intermediary, while R07159 in the acetate phenotype reduces ferredoxin instead. The acetate phenotype further relies on R00229 to transform acetyl-CoA into acetate, which relies on ATP. Meanwhile, R00229-G is responsible for this conversion in the ethanol phenotype. R00229-G uses GTP instead of ATP. These differences are highlighted in Figure 3A, box 2.

Finally, we can visualise the difference in fluxes being directed towards the production of ethanol in each phenotype. Most of the acetate produced in the acetate phenotype is exported into the extracellular space (Acetate_e), while all the acetate in the ethanol phenotype is further metabolised into ethanol, to then be exported (Ethanol_e). This demands increased activity in both AOR_acetaldehyde and R00746. AOR_acetaldehyde converts acetate into acetaldehyde while oxidising ferredoxin and R00746 oxidises NADPH to convert acetaldehyde into ethanol (Figure 3, box 3).

Another key metabolic difference between the acetate and the ethanol phenotypes, that is partially explained by genetic modifications in the ethanol phenotype, is how the system cycles sodium. The acetate phenotype relies on the oxidation of ferredoxin to import sodium through the mbh reaction, which is knocked out in the ethanol phenotype. As an alternative, the ethanol phenotype utilises CODH, an inserted sodium importer that converts intracellular CO into CO2 in the process (Figure 3, box 4).

### Case 3: Identifying mechanistic drivers in the acetate and ethanol phenotypes

Genetic and reaction manipulations are certainly influencing end-metabolite production differences between the acetate and ethanol phenotypes. However, Vailionis *et al.* trialled several similar engineering strategies to improve ethanol production in iPfu, with most of them being unsuccessful (5).

Therefore, we utilised MMINT to identify and show the underlying drivers that make this particular engineering strategy a successful one.

The portions of the networks that we visualise in Case 2 hint at differences in terms of cofactor-cycling between the analysed phenotypes, with non-zero flux reactions that require either ferredoxin or NADP/NADPH. First, two key reactions involved in the cycling of ferredoxin (mbh, sodium importer, Fdred (reduced ferredoxin) to Fdox (oxidised ferredoxin); R07159, GAP to 3PG, Fdox to Fdred) in the acetate phenotype are inactive in the ethanol phenotype (Figure 4A), suggesting a reduced reliance on ferredoxin in the latter. However, ferredoxin is crucial for metabolising pyruvate and CoA into Acetyl-CoA with R01196, a non-zero flux reaction in both phenotypes. It is this need (ferredoxin cycling), and the lack of alternatives, that redirect fluxes from the knocked mbh towards AOR_acetaldehyde, increasing the production of acetaldehyde (Acetaldehyde_c), an intermediary between acetate and ethanol.

Neither the acetate nor the ethanol phenotypes are capable of exporting acetaldehyde; hence, to prevent its accumulation in the intracellular space, the metabolite is further converted into ethanol by R00746, which can be exported to the extracellular space. Yet, this step requires NADPH oxidation. A larger flux in the ethanol phenotype through AOR_acetaldehyde is therefore translated into an increased dependence on NADP/NADPH cycling (Figure 4B). This explains the redirection of fluxes through GAPN (R01058, NADP reduction) in the ethanol phenotype, over R07159 (ferredoxin reduction).

In summary, the phenotypic change between acetate and ethanol production is driven not only by the gene/reaction modifications to the iPfu network, but by the underlying shifts these introduce in terms of cofactors required to maintain redox balance. MMINT visualisation suggests that acetate production in the corresponding phenotype results from its reliance on ferredoxin cycling. Modifications to the reactions in such phenotype result in flux being distributed through reactions that rely both on ferredoxin and NADP/NADPH cycling instead, and lead to the production of ethanol.

## Discussion

This work presents MMINT, a new tool designed to facilitate the exploration and comparison of metabolic flux networks, addressing a key challenge in the field of metabolic modeling. MMINT enables researchers to move beyond static representations of metabolic networks by providing an intuitive, interactive interface for visualizing and interrogating complex flux distributions. As demonstrated through the case studies using the *P. furiosus* GEM, MMINT can illuminate relevant metabolic pathways, highlight differences between phenotypic states, and uncover the underlying mechanistic drivers of these variations. Specifically, our analysis revealed the importance of cofactor balancing in driving the shift from acetate to ethanol production, demonstrating the power of MMINT to identify non-intuitive mechanistic metabolic principles, from which phenotypes emerge.

Genome-scale metabolic modelling is widely used to understand molecular mechanisms underlying metabolic phenotypes. These insights can be leveraged to predict optimal engineering strategies and design microbial strains with desired properties. MMINT enables metabolic discovery by providing a stand-alone GUI to explore flux distributions under alternative metabolic scenarios. MMINT is agnostic to the tool used for flux balance analysis and automatically constructs the full global network *de novo* without reference to an existing map. In contrast to existing tools (e.g. CAVE), MMINT provides novel functionality that allows users to explore and simplify the entire network quicky and easily without specifying a substrate or product. Currently no other visualization or design tools provide the flexibility to explore full CBA solutions and discover underlying metabolic mechanisms with the speed and clarity provided by MMINT.

The case studies presented here highlight the capabilities of MMINT to rapidly explore alternative flux distributions and identify key metabolites and reactions. For example, in the acetate phenotype, MMINT highlights how maltose import and metabolism lead to biomass production, facilitating the discovery of critical pathways by hiding common metabolites. This targeted visualization aids in understanding the specific biochemical routes utilised by different metabolic phenotypes. Case Studies 2 and 3 also demonstrate how comparing flux distributions across metabolic phenotypes with MMINT identified underlying drivers such as cofactor cycling. MMINT visualizations highlight that phenotypic changes between acetate and ethanol production are driven by gene/reaction modifications and shifts in cofactor requirements for redox balance. MMINT suggests that acetate production relies on ferredoxin cycling, while engineering modifications shift fluxes to rely on NADP/NADPH cycling, leading to ethanol production. This suggests that cofactor-driven engineering strategies may successfully enhance ethanol production.

Similar cofactor-driven engineering strategies have been reported in the literature. For example, an alternative NADPH regeneration pathway has been engineered into *Saccharomyces cerevisiae* to improve ethanol production (41). Furthermore, phenolic acid production has been improved by systematically engineering cofactor supply and recycling (36). Thus, MMINT offers a powerful platform for dissecting and comparing metabolic networks, revealing mechanistic insights crucial for understanding and engineering metabolic phenotypes. The integration of cofactor-driven strategies further enhances the potential of metabolic engineering to optimize production processes and achieve desired phenotypic outcomes.

## Data availability

The data underlying this article are available in the article and in its online supplementary material. MINTT is available in https://doi.org/10.6084/m9.figshare.26409328

## Supplementary Data statement

Supplementary Data are available at NAR Online

## Acknowledgements

Not applicable

## Supplementary files

Supplementary file 1

Description: Metabolite and reaction details of base models b971180d_YM_closed_OEadhf and b971180d_YM_GAPN_CODH_closed_OEadhf.

Format: xlsx

Supplementary file 2

Description: Model b971180d_YM_GAPN_CODH_closed_OEadhf with translated metabolites Format: SBML

Supplementary file 3

Description: Flux distributions after FBA in the ethanol phenotype Format: csv

Supplementary file 4

Description: Flux distributions after FBA in the acetate phenotype Format: csv

Supplementary file 5

Description: Supplementary figures and step-by-step guide to replicate networks shown in Case Studies 1 to 3

Format: docx

Supplementary file 6

Description: Python scripts and base models required to reproduce the flux distributions for the acetate and the ethanol phenotypes

Format: zip

