## Supplementary file 5 for "MMINT: a Metabolic Model Interactive Network Tool for the exploration and comparative visualisation of metabolic networks"

### MMINT: [Final Title]

#### Instructions for network visualisation

The present guide provides node and edge legend details (Figure S1) for network elements in MMINT and a step-by-step process for replicating the network visualisations shown in case studies and figures 2, 3 and 4, correspondingly.

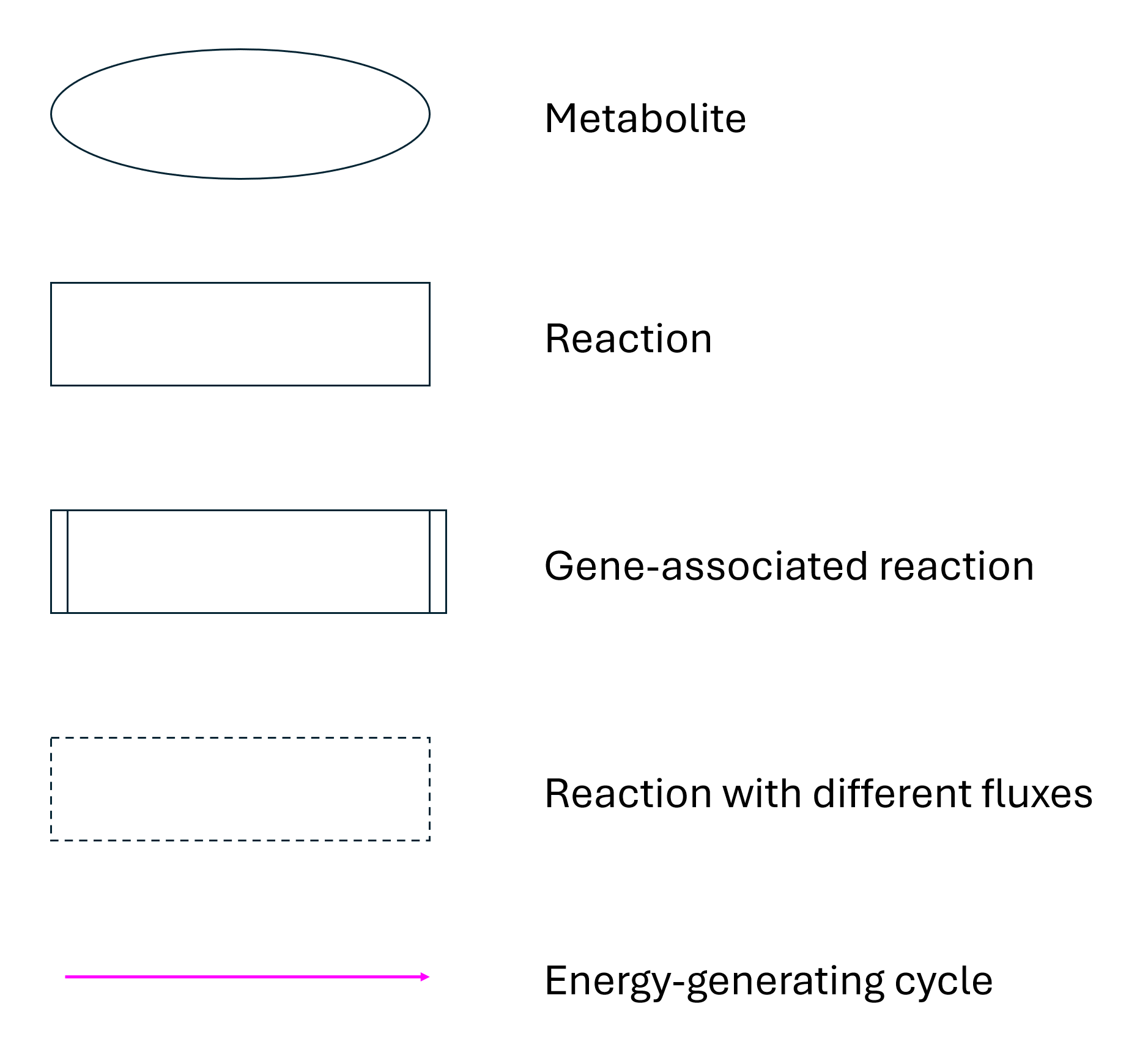

Figure S1 Network elements that can be displayed in MMINT. Colours may vary based on user input.

##### Initial setup

Launch MMINT and load Supplementary file 2 (translated model) in the Data file slot. Load Supplementary file 3 in the Solution file 1 slot to replicate the network from Case Study 1. Load Supplementary file 3 in the Solution file 1 slot and Supplementary file 4 in the Solution file 2 slot to replicate the network from Case Study 2 and 3.

##### Case 1: Targeted network visualisation

To reproduce the network visualisation shown in this section (Figure 2B) in MMINT:

1. Select Maltose_e as the sole Media element
2. Add Biomass_c to Product
   a. Right click on M_Biomass_c on the graph frame
   b. Select Add to Product on the dropdown menu
   c. Select it as the only item in Products
3. Click Apply
4. To hide metabolites in MMINT:
   a. Right click on the metabolite to be hidden on the graph frame
   b. Select Hide on the dropdown menu
5. Hide the following (pseudo)metabolites in the specified order:
   a. Phosphate_c
   b. ADP_c
   c. Traceelements_c
   d. Diphosphate_c
   e. AMP_c

##### Case 2: Solution comparison

To reproduce the network visualisation shown in Figure 3A in the main manuscript:

1. Select Maltose_e as the sole Media element
2. Select Acetate_e and Ethanol_e as Products
3. Click Apply
4. Hide common metabolites/cofactors, including:
   a. Phosphate_c
   b. ADP_c
   c. ATP_c
   d. H+
   e. H2O_c
   f. NADPH_c
   g. NADP_c
   h. Diphosphate_c
   i. GDP_c
   j. Fdox_c (Oxidised ferredoxin)
   k. Fdred_c (Reduced ferredoxin)
5. Trace metabolites from Maltose to the relevant end metabolites by cycling between Explode (reactions and metabolites) – Hide (metabolites) – Unexplode (reactions and metabolites)
   a. To explode reactions and metabolites first right click on the element to be exploded and then select Explode/Unexplode from the dropdown menu.
6. Figure S2 shows in full the reactions and metabolites involved in the conversion of maltose to acetate and/or ethanol in the studied phenotypes. Follow the cycle described in point 5 to add or remove different network elements until the desired outcome has been achieved.

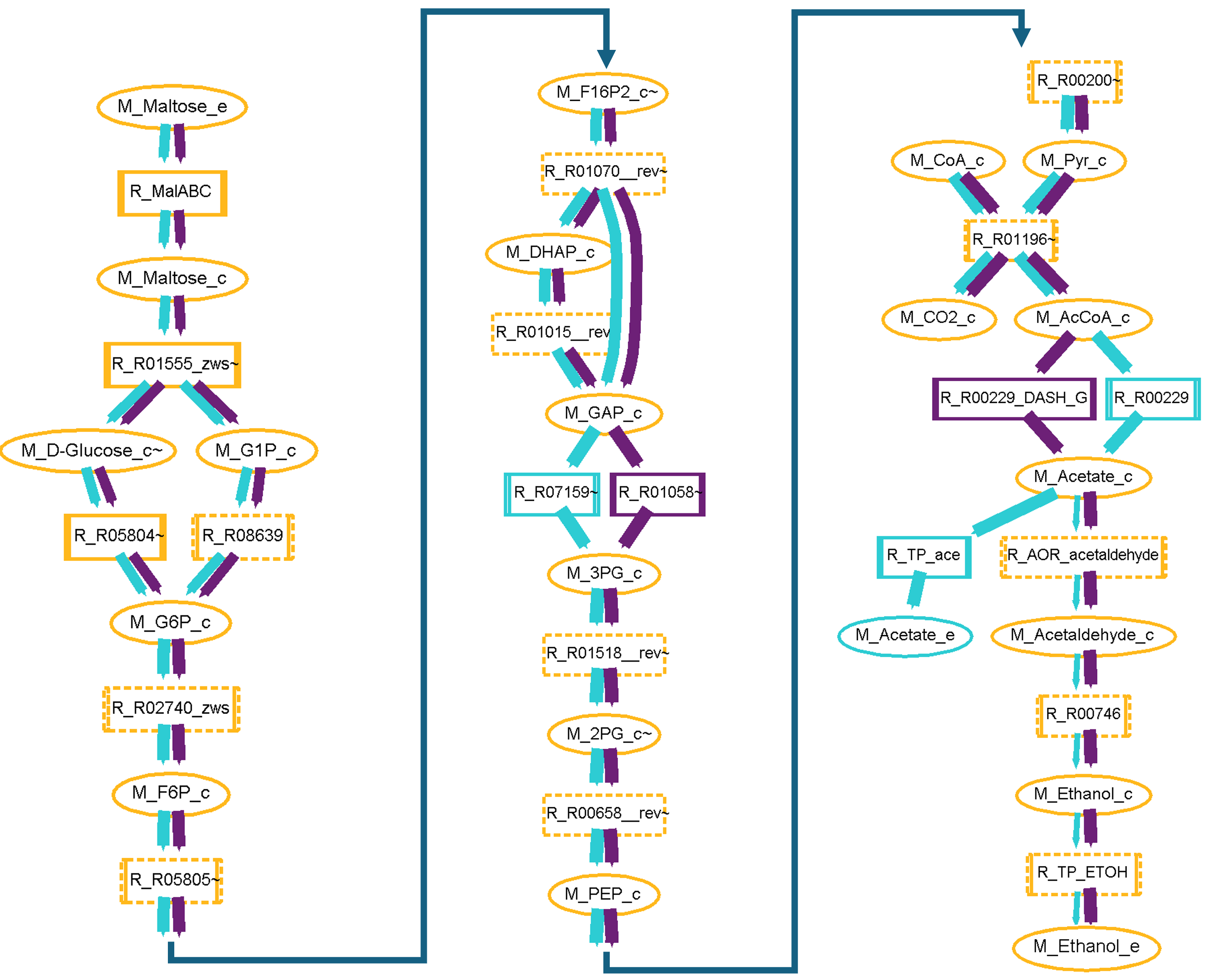

Figure S2 Carbon fluxes from Maltose_e to main metabolites (Acetate_e and Ethanol_e) in the Acetate and Ethanol phenotypes.

To reproduce the network visualisation shown in Figure 3B in the main manuscript:

1. Select CO_e as sole Media metabolite
2. Select CO2_e as sole Product metabolite
3. Explode reaction CODH. MINTT should now show Sodium_e and Sodium_c as network metabolites.
4. Add Sodium_e as Media and Sodium_c as Product. Click Apply.
5. Remove CO_e and CO2_e from Media and Product, respectively. Make sure Fdox and Fdred are not in the list of Hidden metabolites.
6. Explode the following elements in the network:
   a. Sodium_e
   b. Reactions mbh, mrp and atps
7. Optionally, hide metabolites Hydrogen_c and H+_e.

While the steps above will result in a network with the same elements as the one shown in Figure 3B, the spatial distribution of such elements may differ.

##### Case 3: Case 3: Identifying mechanistic drivers in the acetate and ethanol phenotypes

To reproduce the network shown in Figure 4A:

1. Under Algorithm, select Highlight Flood
2. Add Fdred to Media and Fdox to Product
3. Explode both metabolites
4. Identify and explode relevant reactions:
   a. R01196
   b. mbh
   c. AOR_acetaldehyde
   d. R07159
5. Optionally, hide less relevant metabolites
6. Un-explode Fdred and Fdox to hide less relevant reactions

To reproduce the network shown in Figure 4B:

1. Add NADPH to Media and NADP to Product
2. Explode both metabolites
3. Identify and explode relevant reactions:
   a. R00746
   b. R01058
   c. R07181
4. Optionally, hide less relevant metabolites
5. Un-explode NADPH and NADP to hide less relevant reactions
