## Supplementary file 6 for "MMINT: a Metabolic Model Interactive Network Tool for the exploration and comparative visualisation of metabolic networks": Reproduce_paper_solutions.ipynb

```
# %% [code]
import cobra
import pandas as pd

# %% [code]
ethanol_model = cobra.io.read_sbml_model('models/b971180d_YM_GAPN_CODH_closed_OEadhf.sbml')

# %% [code]
def add_fixed_constraint(model, rxn, value):
    """
    model = cobra formatted model
    rxn = string of reaction id
    value = a single int/float to constrain the reaction to, or a list/tuple like (lb, ub)
    """
    try:
        lower = value[0]
        upper = value[1]
    except (TypeError, IndexError):
        lower = value
        upper = value
    constraint = model.problem.Constraint(
        getattr(model.reactions, rxn).flux_expression,
        lb=lower, ub=upper)
    model.add_cons_vars(constraint)
    return constraint


# %% [code]
temp_ethanol_model = ethanol_model.copy()  

with temp_ethanol_model:
    temp_ethanol_model.reactions.TP_ETOH.lower_bound = 3.29265
    temp_ethanol_model.reactions.mbh.knock_out()
    mal_upt = float(cobra.flux_analysis.flux_variability_analysis(temp_ethanol_model, "MalABC").maximum)
    add_fixed_constraint(temp_ethanol_model, "MalABC", mal_upt)
    sol = temp_ethanol_model.optimize()
    acetate_flux = sol.fluxes['EX_C00033_LSQBKT_e_RSQBKT_']
    ethanol_flux = sol.fluxes['EX_C00469_LSQBKT_e_RSQBKT_']
    print('Temp model')
    print(f'Biomass: {sol.objective_value}')
    print(f'Acetate: {acetate_flux}')
    print(f'Ethanol: {ethanol_flux}')
#     ethanol_solution_df = pd.DataFrame(sol.fluxes)
#     ethanol_solution_df.to_csv('ethanol_solution/base_fluxes.csv')

# %% [code]
base_model = cobra.io.read_sbml_model('models/b971180d_YM_closed_OEadhf.sbml')

# %% [code]
base_temp_model = base_model.copy()

with base_temp_model:  
    base_temp_model.reactions.TP_ETOH.lower_bound = 3.29265
    mal_upt = float(cobra.flux_analysis.flux_variability_analysis(base_temp_model, "MalABC").maximum)
    add_fixed_constraint(base_temp_model, "MalABC", mal_upt)
    sol2 = base_temp_model.optimize()
    acetate_flux = sol2.fluxes['EX_C00033_LSQBKT_e_RSQBKT_']
    ethanol_flux = sol2.fluxes['EX_C00469_LSQBKT_e_RSQBKT_']
    print('Temp model')
    print(f'Biomass: {sol2.objective_value}')
    print(f'Acetate: {acetate_flux}')
    print(f'Ethanol: {ethanol_flux}')
    # acetate_solution_df = pd.DataFrame(sol2.fluxes)
    # acetate_solution_df.to_csv('acetate_solution/base_fluxes.csv')
```
